# Gaze following in Archosauria – Alligators and palaeognath birds suggest dinosaur origin of visual perspective taking

**DOI:** 10.1101/2022.09.23.509198

**Authors:** Claudia Zeiträg, Stephan A. Reber, Mathias Osvath

## Abstract

Visual perspective taking marks an evolutionary shift in the formation of advanced social cognition. In humans, it is foundational for our communication and understanding of others. Visual perspective taking has also been found in some other primates, a few songbirds, and some canids. Despite its essential role for social cognition, visual perspective taking has only been fragmentedly studied in animals, leaving its evolution and origins uncharted. To begin to narrow this knowledge gap, we investigated extant archosaurs by comparing the neurocognitively least derived extant birds – palaeognaths – with the closest living relatives of birds, the crocodylians. In a gaze following paradigm, we showed that palaeognaths engage in visual perspective taking and grasp the referentiality of gazes, while crocodylians do not This suggests that visual perspective taking originated in early birds or non-avian dinosaurs – likely earlier than in mammals.

## INTRODUCTION

The advent of visual perspective taking represents a key event in the evolution of social cognition. It marks the transition from a unidirectional to a multidirectional frame of reference in social situations, providing information about the world that would otherwise remain out of reach, and offering new beneficial ways of navigating social environments. Among other things, perspective taking lays the foundation for so-called referential communication, where one refers to a jointly perceived object or event. It also forms the bedrock for ascribing beliefs and mental states to other individuals. However, the most basic form of perspective taking, upon which further skills rely, is the generalization from an egocentric to an allocentric visual viewpoint. Put simply: appreciating that someone else can see what you cannot, and consequently being able to recognize what the other one is attending to. This ability can be identified in the ways humans and other animals follow the gazes of others. Visual perspective taking is revealed in the most advanced form of gaze-following, where the gaze target of the other is blocked from the onlooker’s view, causing the onlooker to reposition itself to see what the other is seeing. The ability to take someone else’s visual perspective in this way, is known as geometrical gaze following [e.g. 1].

Despite its foundational role in social cognition, studies on visual perspective taking have largely lacked a phylogenetic focus, leaving a patchy understanding of cognitive evolution in general. To date, geometrical gaze following has only been found in apes, monkeys, wolves (and dogs), corvids and starlings [2-6]– diverse lineages that all have arisen after the K-Pg boundary, a period witnessing extensive neurocognitive evolution [7]. Hence, we are currently uninformed about one of the major transitions in social cognition. Considering the growing evidence that mammals and birds – separated by 325 million years – have evolved similar cognitive repertoires independently [8], and the fact that geometrical gaze following has only been found in few mammalian and avian species, there are good reasons to assume that visual perspective taking has arisen separately multiple times. It is essential to study each lineage in deep time to better understand the principles of socio-cognitive evolution. Such studies, in combination with research on brain evolution, may shed light on the timing, selective pressures, and possible alleviations of evolutionary constraints.

To begin establishing when visual perspective taking arose in Sauropsida (the lineage of reptiles and birds), we used the paleontological inference method of extant phylogenetic bracketing [9]. By comparing the gaze following repertoire of crocodylians with that of palaeognath birds, we phylogenetically bracketed the dinosaur lineage leading to birds as closely as possible. Crocodylians are the closest living relatives of birds. They have had slow evolutionary rates [10], and seem to have largely retained an ancestral brain morphology [11]. Palaeognath birds, on the other hand, are the most neurocognitively plesiomorphic extant birds, making them in this regard more similar to paravian dinosaurs than any other bird taxa [7, 12].

The study of gaze following has its roots in developmental psychology and comprises an extensive research program, which has been successfully adopted by animal research. Early on, gaze-following was divided into two qualitatively different levels, a high and a low level [13]. The high-level affords the aforementioned geometrical gaze following, while low level gaze-following is an almost reflexive co-orientation with the visual direction of the other [14]. The low level does not require prior expectations to find anything in the gaze direction, or representations of the referentiality of the gaze, but is an adaptive reaction that leads to noticing objects or events that could otherwise have been missed. Such gaze-following is mediated by conserved sub-cortical structures [15, 16]. Low level gaze-following is commonly tested through gaze-following into the distance experiments, where a demonstrator is lured to gaze either up or to the side. An onlooker capable of this skill is expected to co-orient with the gaze direction of the demonstrator. Low level gaze-following develops far earlier in children than high level gaze following, with an onset between 3 and 6 months of age [e.g. 17, 18]. Gaze-following into the distance has so far been found in all studied amniotes, ranging from mammals to birds and reptiles [e.g. 19, 20, 21].

As mentioned, high level gaze-following, on the other hand, is a notably more advanced form. It presupposes expectations of finding something in the other’s line of gaze, and that this gaze reference can only be found if one changes one’s own perspective. This is the reason it is tested in the geometrical gaze following paradigm, with barriers blocking the view, that must be circumvented. Unsurprisingly, this form of gaze following is suggested to be mediated by various cortical areas [22];although the avian homologues for such gaze-following still need to be determined. In children, geometrical gaze-following is not seen until the age of 18 months [17].

Another central gaze-following behaviour, that thus far has only been reported in humans, apes, and Old World monkeys [e.g. 2, 23, 24] is the so-called “checking back”. “Checking back”-behaviour is instigated when no object of interest is identified in the other’s line of gaze, or when the gaze direction and its target are incongruent. The observer will then look back at the other in an apparent attempt of re-tracking the gaze direction. The “checking back”-behaviour is regarded an essential diagnostic behaviour for the onlooker’s representation of the referentiality of the other’s gaze, i.e. that it is pointing towards something [25]. “Checking back” thereby reveals a violation of the expectancy to find a gaze target in the observed gaze direction.

Visual perspective taking, as displayed in geometrical gaze following, does not imply the representation of others’ epistemic or perceptual states. Rather, it is a form of functional representation, leading to behaviors that correspond to the fact that the other has a different perspective and that its gaze refers to an object.

Furthermore, visual perspective taking is traditionally divided into a level I and II [26]. Level I enables taking into account *what* (or that “something”) lies in the line of gaze of the other, or in other words, what the other can or cannot perceive. In children this level develops between 18 and 24 months [e.g. 27, 28, 29]. Level II, on the other hand, requires the adoption of the spatial viewpoint of the other, and hence taking into account *how* the world is perceived from that perspective. One understands that the same thing oneself sees, is perceived differently from the angle of the other. This is considerably more advanced, and does not develop in children until the age of 4-5 years of age [e.g. 30]. It has been suggested that while geometrical gaze following cannot reveal level II perspective taking, it forms the embodied pre-cursor to develop or evolve it. Repositioning the body provides an experience of the other’s perspective, which in turn can be used in mental simulations of one’s own body positions to understand others [31]. Taken together: geometrical gaze-following is a sophisticated embodied sensory-motor process that anchors the most advanced forms of social cognition.

To investigate potential level I visual perspective taking skills in extant archosaurs, which phylogenetically bracket the extinct Dinosauria, we tested 30 individuals from five archosaur species (six per species) for their ability to follow conspecific gaze: emus (*Dromaius novaehollandiae*), greater rheas (*Rhea americana*), elegant crested tinamous (*Eudromia elegans*), red junglefowl (*Gallus gallus*), and American alligators (*Alligator mississippiensis*). The three palaeognath species represent different phylogenetic nodes within that group, and different socio-ecologies, as well as flightlessness and volant flight [e.g. 32]. The red junglefowl were added as an outgroup of basal neognaths, belonging to the lineage Galloanserae that diverged from Neoaves (the other large group of neognaths) before the K-Pg extinction event. The animals were tested in three gaze-following experiments: following gaze into the distance

(1) up and (2) to the side, and geometrically (3) behind a barrier (for experimental setups see Figure 1). The potential presence of “checking-back” behaviour was studied in all three experiments.

**Figure 1:**
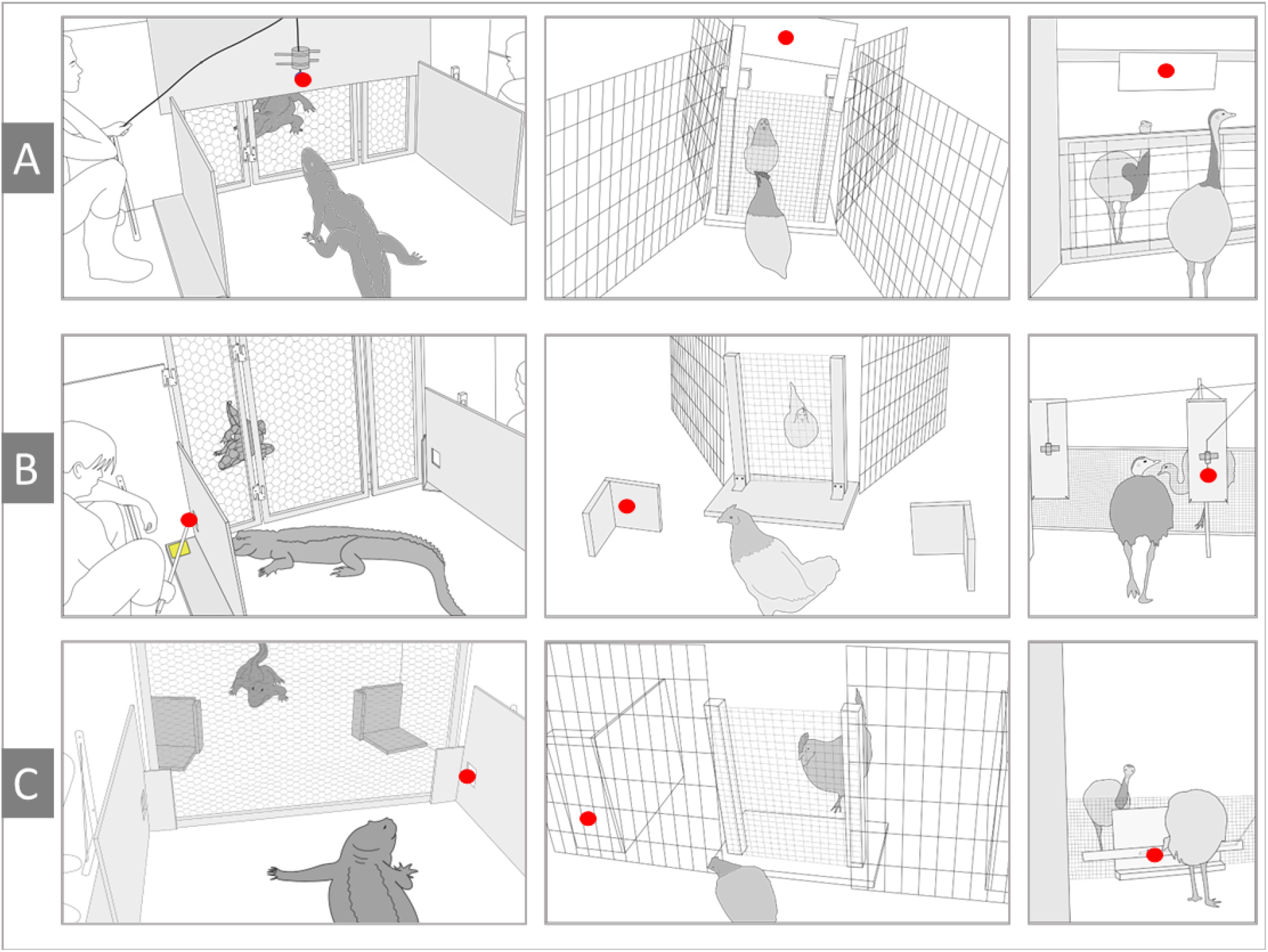
All experimental setups (from left to right) for alligators, small birds (red junglefowl and elegant crested tinamous), and large birds (emus and rheas). Row A) Setups for Experiment 1 (gazing up). Row B) Setups for Experiment 2 (gazing to the side). Row C) Setups Experiment 3 (geometrical). Red dots depict stimuli used to lure demonstrators’ gazes (for more information about stimuli see Supplementary Material).

## RESULTS

### Gaze-following into the distance and geometrical gaze-following

All tested species followed conspecific gazes into the distance. In Experiment 1 (gazing up), all birds performed at a comparable level (see Figure 2). However, the alligators did not respond by looking up, but instead turned around and looked behind themselves at a significant level (see Figure 2).

**Figure 2:**
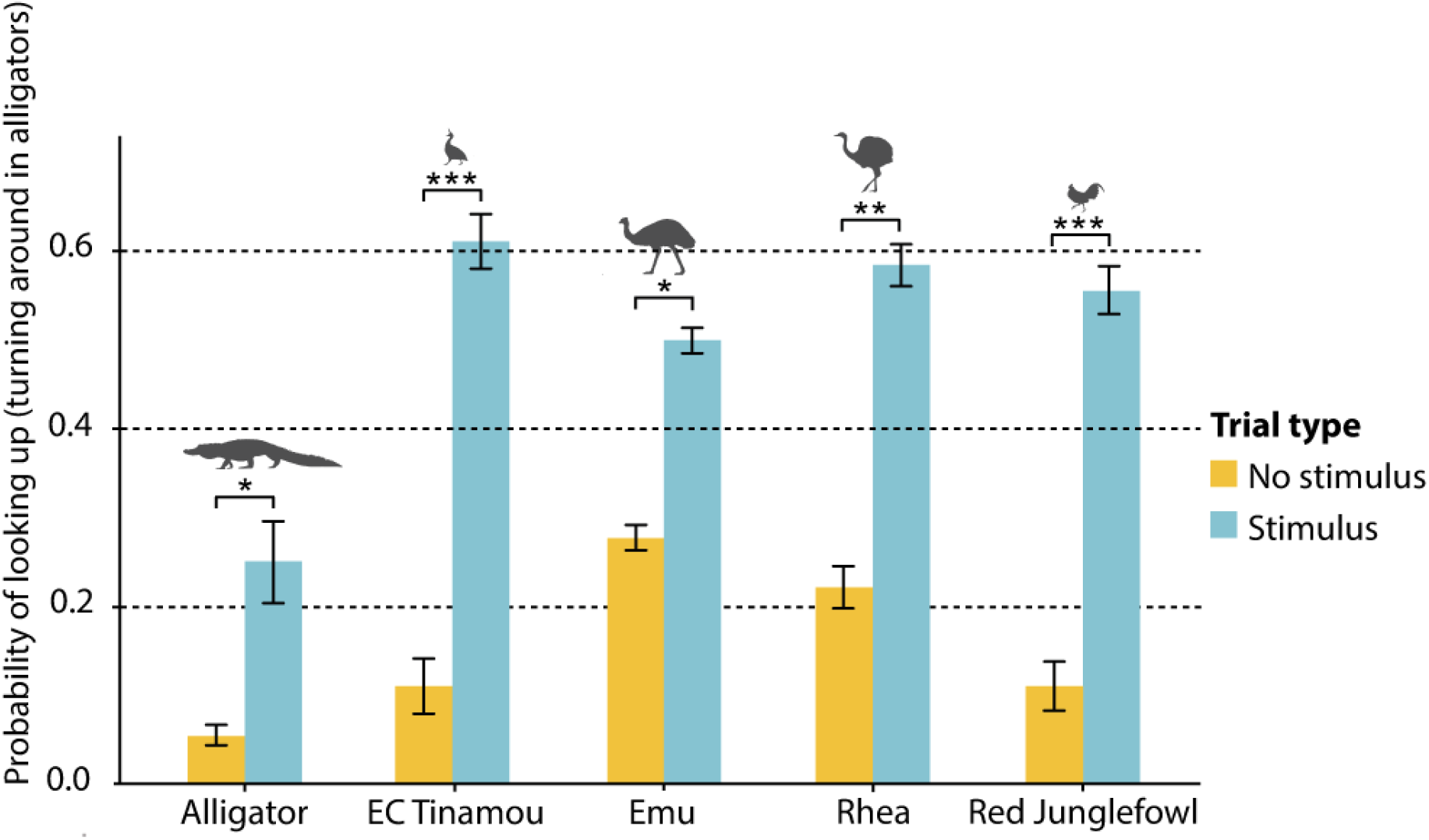
Gaze-following into the distance: Up. Probability of looking up (turning around in alligators) in demonstrator condition of Experiment 1. All bird species looked up significantly more often in trials with a stimulus present (a demonstrator gazing up) compared to trials with no stimulus (likelihood ratio test, χ^2^ >4.55, df = 1, p<0.033). Alligators reacted by turning around and looking behind themselves. They did so significantly more often in trials with a stimulus present (likelihood ratio test, χ^2^ = 5.77, df = 1, p = 0.016).

In Experiment 2 (gazing to the side), all birds passed the test at similar rates. The alligators also passed, but with a notably lower rate than any bird species (see Fig 3).

**Figure 3:**
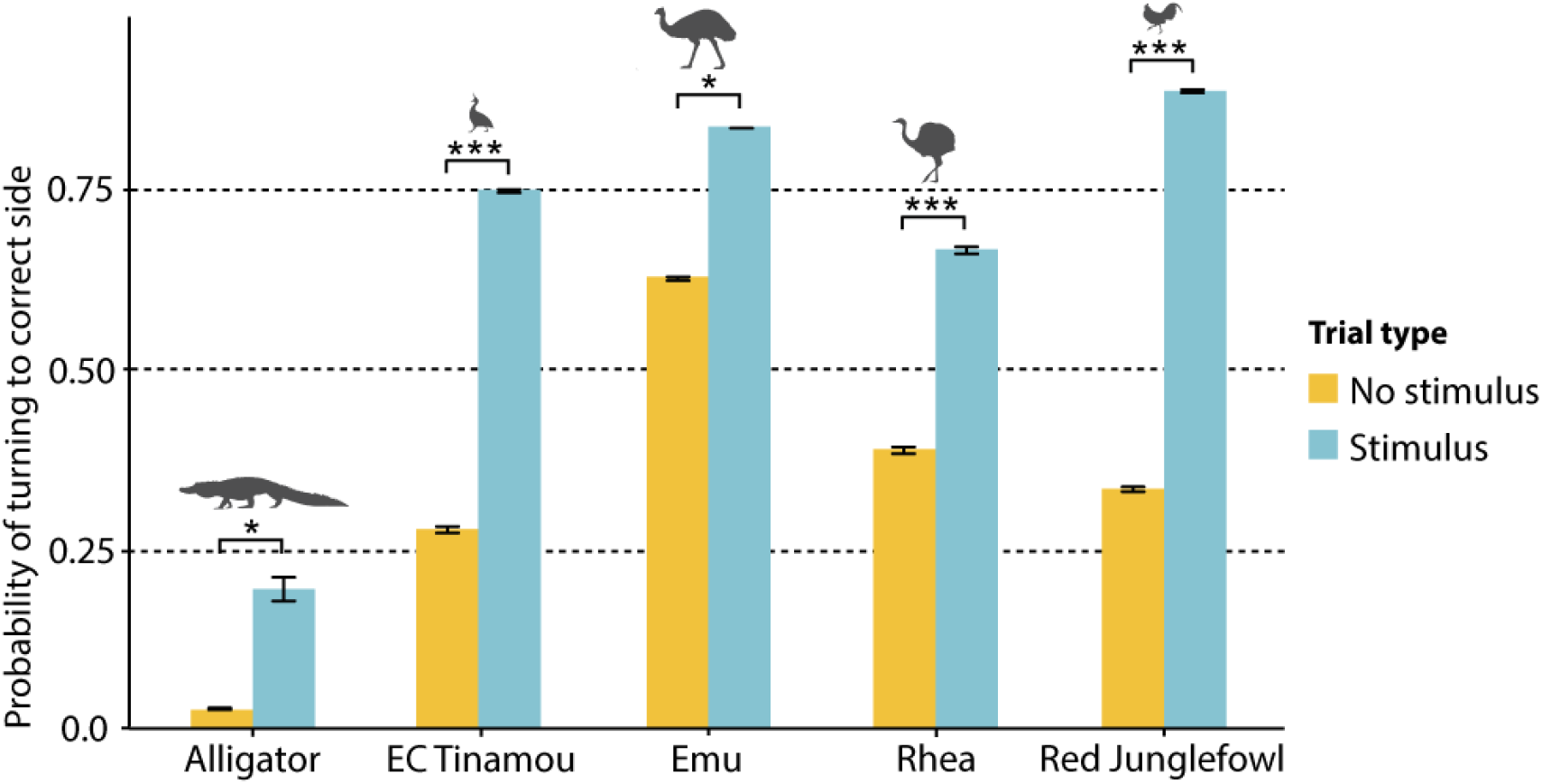
Gaze-following into the distance: Sideways. Probability of turning to correct side in demonstrator condition of Experiment 2. All bird species turned more to the correct side in trials with a stimulus present (likelihood ratio test, χ^2^ >3.88, df = 1, p<0.049). No significant difference in gaze-following rate between bird species was found. Alligators followed gaze at significantly lower rates compared to birds (likelihood ratio test, χ^2^ = 15.055, df = 4, p = 0.0046). EC Tinamou = elegant-crested tinamou.

There is a clear difference in the frequency of gaze follows into the distance (Experiment 1 and 2) between alligators and birds (see Figure 4), even when regarding the turning around behaviour by the alligators in Experiment 1 as a gaze-following response. There is no significant difference between the different bird species.

**Figure 4:**
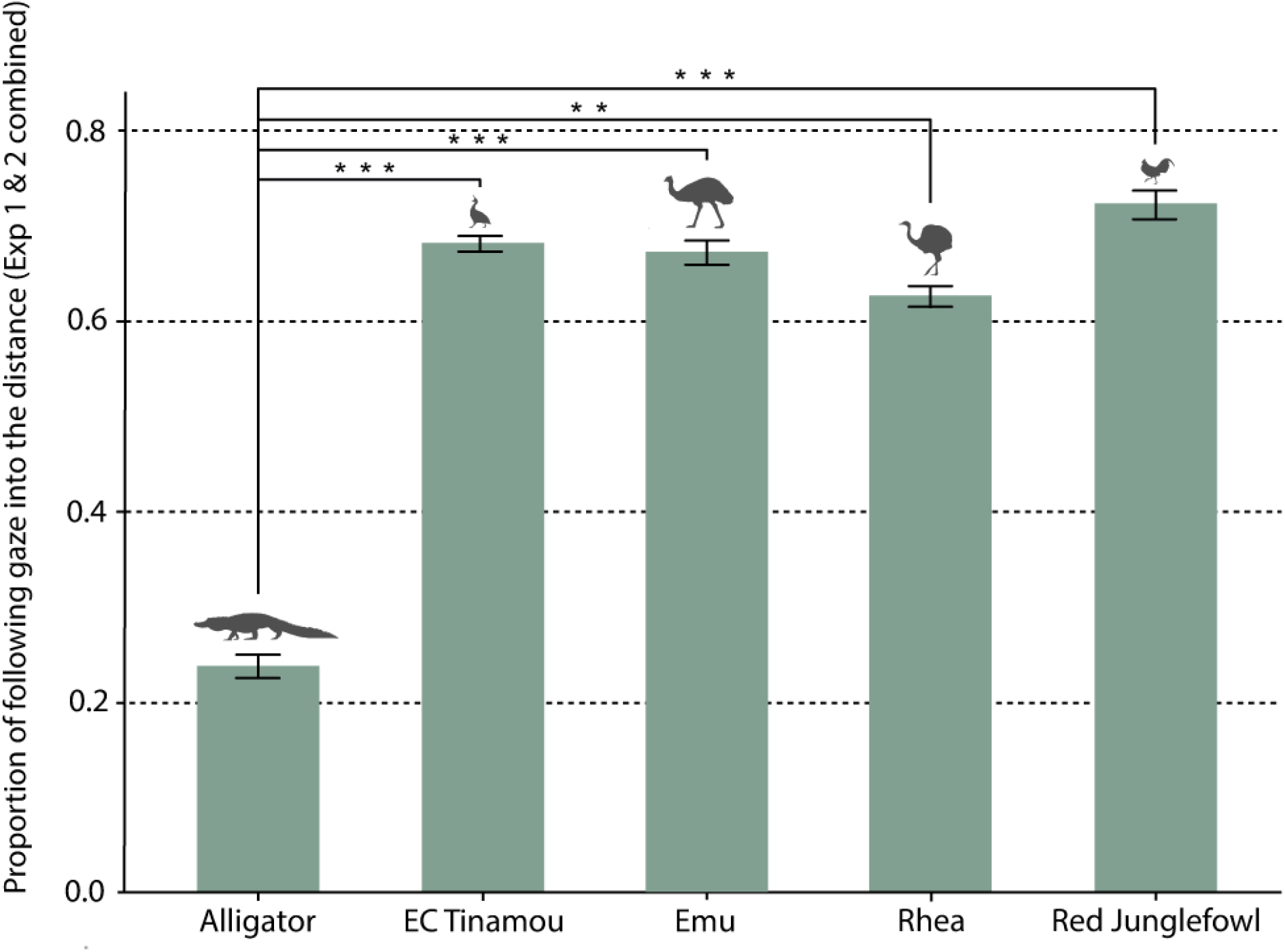
Proportion of gaze-following into the distance. Species had a significant effect on probability of gaze-following (likelihood ratio test, χ^2^ = 26.407, df = 4, p<0.001). Gaze-following proportions were significantly higher for birds (elegant-crested tinamou; in this graph “EC Tinamou”: 0.68, emu: 0.67, rhea: 0.63, red junglefowl: 0.72) compared to alligators (0.24).

All bird species followed gaze geometrically, and at comparable rates (see Figure 5). The alligators, however, did not reveal any geometrical gaze-following in the test.

**Figure 5:**
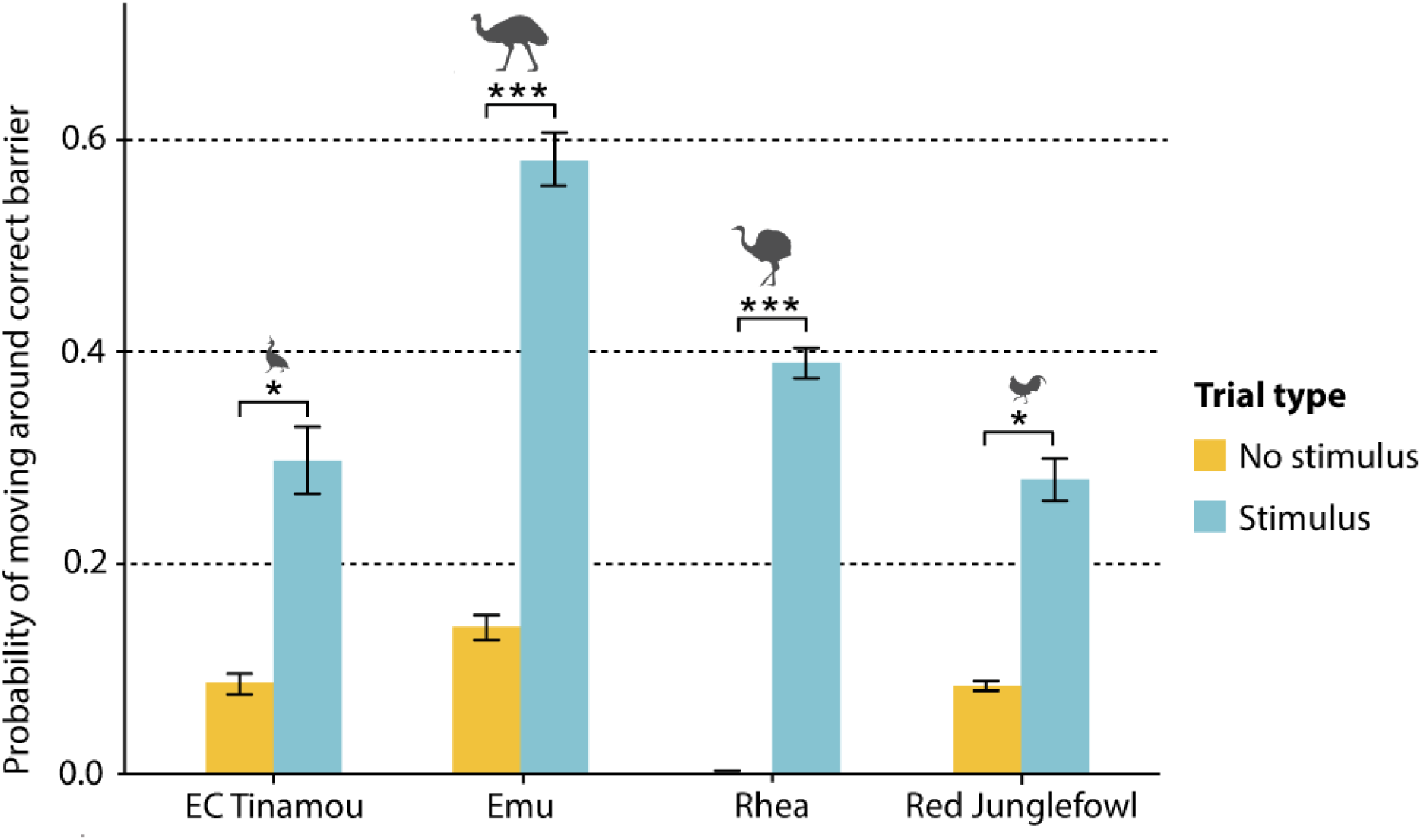
Geometrical gaze-following. Probability of moving around correct barrier in demonstrator condition of bird species (Experiment 3). Alligators did not follow gaze geometrically. Between birds, no significant effect of species was found, but there is a trend for a higher proportion in emus (Z= 1.93, p=0.054). All bird species moved around the correct barrier significantly more often in trials with a stimulus compared to trials without a stimulus (likelihood ratio test, χ^2^ = 33.74, df = 1, p<0.001). EC Tinamou = elegant crested tinamou.

### “Checking-back” behaviour

All bird species engaged in “checking-back” behaviour, but the alligators did not. There was a significant species effect among the birds on the probability of checking back in Experiment 3, the geometrical gaze-following (likelihood ratio test: χ^2^ = 9.73, df = 3, p = 0.021). This difference, however, is likely caused by differences in the experimental setups because of varying body sizes. While the larger birds (emus and rheas) could effortlessly check back by lifting their head, the smaller birds (junglefowl and tinamous) had to walk out from behind the barrier to be able to see the demonstrator again, as they were too small to look over it. This is likely the reason why larger birds have been found to check back more often in the geometrical experiment, while in the other two experiments, all birds checked back at comparable rates. This is evident from the fact that this difference was found *after* relocation, i.e. after birds had looked behind the barrier.

“Checking-back” behaviour will lead to a renewed gaze follow towards the target if the demonstrator is still looking at it. In 27% of the “checking-back” instances, the observer again followed the gaze towards the target. In these instances, the demonstrators’ gazes lasted on average 30% (0.73 seconds) longer, indicating that the demonstrator was still gazing towards the target.

## DISCUSSION

This is the first study on visual perspective taking in palaeognath birds and crocodylians. The palaeognath birds, and the junglefowl, show a gaze-following repertoire on par with apes and some Old World monkeys, including behaviors diagnostic of the expectation of a gaze reference. The alligators’ performance is mostly similar to other non-avian reptiles and appears to be restricted to the low-level form of gaze-following into the distance. Collectively, this suggests that visual perspective taking, along with representations of the referentiality of gazes, originated in Dinosauria. It is also likely that these cognitive skills arose earlier in this lineage than in Mammalia, perhaps due to a sophisticated diurnal vision that yet had to evolve in mammals.

Apart from the current study, only one reptile species – the central bearded dragon (*Pogona vitticeps*) – has been tested for geometrical gaze-following [33]. Just as alligators, they did not exhibit these high-level gaze-following skills. However, all previously studied reptiles follow gaze into the distance [21, 33, 34]. This indicates that low level gaze-following skills are shared among reptiles, but that visual perspective taking might be absent, suggesting a comparable repertoire in ancestral archosaurs.

Interestingly, however, the alligators do not follow gazes upwards, but instead turn around. This contrasts with all tested terrestrial non-avian reptiles, which co-orient with upward gazes [21, 33, 34]. It may reflect crocodylians’ adaptation to a life at the water surface, which is apparent in the horizontal arrangement of their sensory organs, and retinal ganglion cells in the eye [35]. Perhaps they mainly raise the head to see further ahead over the surface, rather than up, which would then be at a location behind and not above the observer. Turning around would then entail gaze-following outside one’s own field of vision, which is a form of geometrical gaze-following. Another interpretation is that turning around is an appeasing response, as snout lifting is a submissive signal [36], however such a response has never been reported, nor observed by us in any other situation. The turning around is likely a response to gaze, however, as alligators show no geometrical gaze-following in Experiment 3, it could be a taxon-specific response due to its potential adaptive importance at the water surface, or it could represent an evolutionary early form of geometrical gaze-following.

That geometrical gaze-following was shown by all bird species in our study, indicates that it should be within the repertoire of all birds, given that the species studied represent some of the neurocognitively most conserved taxa. Previously, geometrical gaze-following in birds has only been identified in two corvid species, common ravens (*Corvus corax*) and rooks (*Corvus frugilegus*) [5, 37] and in one other songbird, the European starling (*Sturnus vulgaris*) [6]. On the other hand, only one other study on birds has investigated geometrical gaze-following. A study on the Northern bald ibis (*Geronticus eremita*), did not find such gaze-following, which would counter our prediction. However, the results probably reflect methodological limitations. Among other things, and in contrast to most studies (including the current one), the animals were not facing each other in the geometrical condition, but stood next to one another which might have distorted the observers prediction of the demonstrator’s visual perspective. The authors themselves also cautioned against the results, and suggested tests with different methods. The best predicition is still that most birds, from all taxa, have this seemingly conserved ability.

“Checking-back” behavior, which was found in all birds, has not been reported outside apes and Old World monkeys. However, our findings suggests that “checking back” is a more widespread behavior than previously thought. It has simply never been described or looked for in other species. The only negative results on “checking back” stem from two species of New World monkey: black-handed spider monkeys (*Ateles geoffroyi*) and tufted capuchin monkeys (*Cebus apella*) [38].

Arguably, “checking back”-behaviors should be within the repertoire of species capable of geometrical gaze-following, as such gaze-following presupposes the expectancy that the other’s gaze is directed at something, which cannot currently be seen. “Checking back” is a behavior signifying such an expectancy. The behavior develops earlier in children than the ability to follow gaze geometrically – 8 months versus 18 months [17, 39]. Indicating that the ability to expect a reference of the gaze, not only precedes, but is a prerequisite for, visual perspective taking. The negative results in the study on New World monkeys, may be experimental artefacts, something the authors also suggested. Indeed, one individual spider monkey in the study was found “checking back” multiple times.

As mentioned, alligators and birds differed in that the alligators did not reveal visual perspective taking (barring the curious turning around behavior) or any “checking back”-behavior. But they also differed in another important measure: the sensitivity to the other’s gaze, which is seen in the proportions of gaze follows in Experiment 1 and 2 (Figures 1 and 2). The birds, on the other hand, had a proportion of gaze follows similar to great apes [e.g.40].

### The potential role of differences in cerebellar size

A major neuroanatomical difference between crocodylians and avians is the radically higher density of neurons in birds, leading to much greater neuron numbers in their brains. The main proportional increase of neurons in the evolution from stem archosaurs to birds is found in the cerebellum [7]. For example, an emu has 20.5 times as many neurons in the cerebellum as a Nile crocodile (*Crocodylus niloticus*), while having 15.75 times as many neurons in the telencephalon. We suggest that the vastly expanded cerebellum provides insights into why the birds, but not the crocodylians (or other reptiles), show visual perspective taking with its accompanying representations of gaze reference. While a large cerebellum is not sufficient for advanced gaze-following, it is most likely a prerequisite.

The cerebellum primarily guides motor control, but is involved in a variety of cognitive processes [41]. This structure is organized in parallel loops through which it simultaneously receives input and sends projections to cortical areas [42]. The highly regular cytoarchitecture suggests a unified mechanism underlying its various functions [43]. An influential theoretical framework proposed for this unifying mechanism is that of the so-called internal forward models.

Such models are top-down processes using prior, instead of immediate, information to guide behavior and to predict behaviors of others [44-47]. Well-developed sensory-motor predictions allow rapid appropriate actions and update quickly when the model does not match the world. This considerably speeds up behavior, as compared to a system that instead continuously responds only to the feedback from the external world (bottom-up).

We propose that the differences in the gaze-following repertoires of alligators and birds is partly explained by more robust internal forward models in birds. Gaze-following is mediated by top-down processes in various action predictions of others [e.g. 48, 49]. The act of gazing can induce the prediction in the observer that the other’s gaze points to “something”. The “checking-back” behavior clearly shows when such expectations are violated, but also that the system is tuned to updating, which is a hallmark of internal forward models [e.g. 47]. The evolution and development of visual perspective taking, and representing referentiality, is likely an embodied process starting out from building sensory-motor forward models of one’s own behavior, which gets extended to other’s basic behaviors [50]. Obviously, more robust internal forward models in the cerebellum, making more detailed and fine-grained predictions, will only arise in the presence of well-developed sensory-motors areas in the pallium (or cortex) which they project on, which is something birds have as compared to reptiles. However, we do not know to what extent the enlargement of the cerebellum seen in the maniraptoran theropods [51], reflects the existence of other brain areas involved in visual perspective taking.

### The origins of visual perspective taking and further research

Palaeognaths are the best available extant neurocognitive models of non-avian – but closely related – paravian dinosaurs, such as dromaeosaurids and troodontids. There are of course differences between the least derived (extant) avian brain and that of extinct non-avian paravians. For example, the presence of the wulst (hyperpallium), and ventrally deflected optical lobes in birds [51], which likely mainly represent adaptations to the visuo-motor requirements of flight. Nevertheless, the palaeognath brain is strikingly more similar to a non-avian paravian brain, than to that of a crocodylian. For example, in size, shape, and proportions between areas [e.g. 7, 12, 52]. But also in the relationship between body and brain size, where palaeognaths fall within the scaling relationship of non-avian paravians, unlike most other birds [53]. One of the central questions, however, is whether the neuronal density was similar between paravians and palaeognaths, because the number of neurons is currently one of the best neurobiological correlates to cognitive performance [e.g. 54, 55]. Palaeognaths have the least derived scaling relationship of neuronal numbers among birds (shared with some neognath taxa), but that still allows about twice as many neurons per volume unit than a non-primate mammal [7, 12]. It has recently been shown that endothermy is highly associated with the extreme increase of neuron numbers [7]. Accumulating evidence from different methodological sources suggests endothermy in at least non-avian paravians [56-58]. There are hence reasons to assume that these dinosaurs had neuronal densities more similar to palaeognaths than to extant reptiles.

Despite the lack of studies on structures in the avian brain corresponding to those in mammals that mediate geometrical gaze-following, it may be the case that they existed in non-avian paravians too, given several similarities to palaeognaths. If so, visual perspective taking could have arisen in the non-avian paravians (or perhaps earlier) and may thus have been present by the Middle Jurassic (ca. 174-163 million years ago). However, if the avian unique wulst, which is an area of visual and somato-sensory integration [e.g. 59, 60], prove central for visual perspective taking, then one would expect that its origin occurred later. There is still no consensus, based on the fossil record, when the wulst appeared, but since it exists in both palaeognaths and neognaths, which according to molecular analyses diverged in Early Cretaceous (about 110 million years ago) [e.g. 32], it should at least have been present then. There indeed exist projections between the wulst and the cerebellum [61]. However, the visual and somatosensory requirements of flight likely exceed those of terrestrial mammals and might therefore represent levels of sensory-motor models beyond what is needed for modelling other’s occluded lines of gaze. Only further research on brain anatomy and brain function in birds, as well as on brain anatomy in extinct dinosaurs (including avians), will help to better pinpoint the origins of visual perspective taking in dinosaurs.

However, as hinted at earlier, the current evidence from mammalian geometrical gaze-following places the skill in lineages that diverged after the K-Pg boundary: Simiiformes (monkeys and apes) and Canidae (where it is only shown in wolves and dogs). That puts the origin of visual perspective taking considerably later in mammals than in birds – with several tens of millions of years. However, if it was not convergently evolved in simians and canids, it should be found in many taxa that diverged since the split of the common ancestor of Primates and Carnivora, ranging from rodents to bats and a long range of others, and its origins would then be traced well before the K-Pg boundary [62] (but still probably many millions of years after its origins in dinosaurs). More gaze-following studies on mammals are needed to provide better understanding, and to disentangle to what extent these skills evolve convergently within mammals.

It is not surprising if visual perspective taking, with accompanying representations of gazes’ referentiality, evolved earlier in dinosaurs than in mammals. The major increase of neurons, which is seen in both mammals and birds – likely as a response to endothermy – might be a prerequisite, but an acute vision may be of additional importance. The benefits of gaze-following are likely enhanced by an advanced visual system, where foveae and color vision seem particularly useful, both of which most likely existed already in non-avian dinosaurs as it exists in reptiles and birds (if not lost due to nocturnal adaptations). Following the gaze of someone who can attend to more details in the environment, as well as see further into it, provides more information, given that the gaze follower itself has similar visual capabilities. Mammals were initially, and for a very long time, primarily nocturnal, and vision had less utility than e.g. olfaction [63]. The most well-developed gaze-following repertoires in mammals are found in simians, and particularly apes. Primates have readapted their vision to diurnal conditions and regained both foveae and color vision. The refinement of the visual system co-evolved with the relative expansion of the primate cerebellum [64], which proportionally increased even more in great apes [65]. Arguably, this expansion led to improved visuo-motor internal forward models for prediction of other’s behaviors, perhaps making apes similar to birds in this regard. Studies on other mammals are needed to understand the role of visual acuity for visual perspective taking, and whether differences may lead to convergent evolution of this skill within mammals. But also, more studies are needed on to what degree other sensory modalities aid in various forms of perspective taking.

Geometrical gaze-following reveals only the most basic forms of visual perspective taking (level I) and cannot attest to more advanced socio-cognitive skills. Decades of research into animal cognition have focused on various aspects of “mindreading”-abilities. Animals’ *mental* perspective taking, such as representations of others’ epistemic states, intentions, desires, or other motivational states, has been intensely studied, where apes and corvids show the highest proficiency (Krupenye and Call, 2019). However, much more research is needed on neurocognitively plesiomorphic animals to better understand the evolution of social cognition.

## MATERIALS AND METHODS

### Experimental Design

We tested 30 subjects of five archosaur species (six per species, for more information on subjects see Supplementary Material) for their ability to follow gazes of conspecifics in three experiments. Testing took place between January 2019 and November 2020. Experiments 1 and 2 tested for gaze following into the distance upwards (Experiment 1) and to the side (Experiment 2). Experiment 3 investigated geometrical gaze following, i.e. tracking gaze around a barrier. Due to limited sample sizes, some individuals of each species were used as both demonstrator and subject. Those individuals first finished all demonstrations before serving as subject to minimize the number of potentially biased trials. Demonstrators were selected based on highest responsiveness to gazing stimuli (described below).

Due to the physical differences of the tested species, three different experimental setups were used within each experiment to create optimal testing conditions. Alligators, large birds (emus and rheas) and small birds (elegant-crested tinamous and red junglefowl), received their own setups, respectively.

A gazing stimulus was used to evoke gazing responses of demonstrators. Demonstrator birds from all bird species besides red junglefowl spontaneously reacted by looking towards the red beam of a laser pointer. Large demonstrator birds (emus and rheas) quickly habituated to this stimulus, so in Experiment 2 and 3 they were lured by pulling a string to release food from an opaque tube. Demonstrators of red junglefowl were conditioned to turn towards the beam of a laser pointer in training sessions prior to the experiments. Demonstrators of alligators were conditioned to turn towards a blue rubber ball. Conditioning was achieved through clicker-training in both species. However, no clicker was used during testing.

Every experiment was divided into two conditions: Demonstrator and no-demonstrator. In the demonstrator condition, subject and demonstrator were present, while only the subject was present in the no-demonstrator condition. Half of the subjects started with the demonstrator condition, the other half with the no-demonstrator condition. Each condition was further divided into two trial types: stimulus and no-stimulus. In stimulus trials, the gazing stimulus was presented, whereas no stimulus was shown in no-stimulus trials. Trial types were pseudorandomized.

Every condition (demonstrator or no-demonstrator) consisted of 12 trials, 6 of each trial type. In stimulus trials of the demonstrator condition, the stimulus was presented until a gazing response of the demonstrator was evoked. In no-stimulus trials of the demonstrator condition, no stimulus was presented, so that the demonstrator was simply present. These trials served as controls for social enhancement.

In stimulus trials of the no-demonstrator condition, only the subject was present while the stimulus was presented for 5 seconds. This served to control if the stimulus was visible from the subject side. In no-stimulus trials of the no-demonstrator condition, no stimulus was shown while only the subject was present. This was done to maintain the same procedure and session length as in the demonstrator condition. In both conditions, the trial lasted for 10 seconds after demonstration (either the demonstrator gazing, or the stimulus being presented without demonstrator present). Only in Experiment 3, alligators were given 1 minute due to the potential amount of walking in this setup.

If a significant difference in orienting responses could be identified between stimulus and no-stimulus trials in the demonstrator, but not the no-demonstrator condition, this difference was most likely caused by the gaze cue of the demonstrator. All trials were videotaped using two JVC EverioR cameras, one behind the subject, and one facing the subject to ensure optimal angles of the heads and eyes of subjects.

### Experimental Setups

#### Experiment 1: Up

In Experiment 1, an opaque screen was mounted on top of the divider that was placed between subject and demonstrator. For alligators, the blue rubber ball that demonstrators were conditioned to turn towards could be lowered into view with a string from an opaque tube attached to this screen on the side facing the demonstrator. For all bird species, the beam of a laser pointer was projected onto the screen on the demonstrator side.

#### Experiment 2: Side

For alligators, two experimenters seated behind 60-centimeter-high wooden barriers on either side of the demonstrator each had a blue rubber ball mounted on a wooden stick. In stimulus trials, one experimenter presented the ball through a cut-out in the wooden barrier they were seated behind. A small wooden barrier in front of this cut-out prevented the subject from seeing the ball. Sides were counterbalanced; each subject received the same number of trials on either side. A sponge underneath the cut-out ensured that no sounds were made when lowering the balls after presentation.

For small birds, two wooden barriers were placed on the demonstrator side on which the beam of the laser pointer could be presented towards the demonstrator.

For large birds, this experiment was conducted in two different ways due to structural differences in the enclosures. For emus, two tall wooden boards were propped up on both ends of the mesh divider on the demonstrator side. Two experimenters stood behind these boards. Each of them held a grape on a stick, which could be shown in a cut-out to lure the gaze of the demonstrator (similar to the alligator setup). For rheas, two wooden boards were hung from poles on each end of the mesh divider. On the side facing the demonstrator, an opaque tube was attached to both boards from which grapes could be lowered into view on a string.

#### Experiment 3: Geometrical

Alligators were exposed to the same setup as in Experiment 2 (side). However, this time, two wooden barriers were placed approximately one meter in front of the mesh barrier on the subject side. The stick with the target ball was in this condition not only shown in the cut-outs but stuck out of them to make the demonstrator gaze behind one of the two barriers on the subject side. In the presence of geometrical gaze following, the subject would have to walk up to the barriers and turn around the indicated one. The barriers were slightly angled, which prevented the subject from seeing behind both barriers simultaneously when placing itself between them.

For small birds, two barriers were placed on the subject side. The beam of the laser pointer was directed to the back of the barrier, so that it was only visible to the demonstrator. In this way, an orientation of the demonstrator towards that barrier looked to the subject as if the demonstrator was looking behind that barrier.

For large birds, a wooden barrier was placed between demonstrator and subject. On the demonstrator side, a contraption was installed on ground level from which a grape could be released from an opaque tube by pulling a string. By showing the grape, the gaze of the demonstrator was lured towards the ground behind the barrier. A successful subject would be expected to lean over the barrier to identify the gaze target.

### Coding Definitions

All videos were coded using Solomon Coder [66]. When coding trials of all three experiments, we coded “target location” and “checking back”. “Target location” had different definitions depending on the experiment, but generally referred to the location where the gazing stimulus was shown. In Experiment 1, the target location was the panel above the divider. We coded “target location” every time a subject looked up towards that panel. For alligators, we additionally coded “turning around”, which was defined as the subject turning more than 90° away from its initial position. In Experiment 2, the target location was the side where the stimulus was shown, or the side the demonstrator looked towards. In no-stimulus trials, we pre-determined “correct” sides randomly and coded “target location” if the subject turned towards that side. We only scored first orientations of subjects in this experiment. The same method was applied to Experiment 3 of the small birds and alligators. Experiment 3 of the large birds did not include sides, but only had one “target location”, the ground behind the barrier. “Target location” was only coded when subjects relocated themselves around barriers (or looked over the barrier in large birds) and not when they looked towards that location. Additionally, we coded the latency of “target location” for each experiment, either from trial onset in no-stimulus trials, or from the onset of the stimulus (the gazing stimulus in the no-demonstrator condition, or the gaze of a demonstrator in the demonstrator condition). We coded “checking back” when a subject looked towards the target location and then back at the demonstrator. We moreover recorded whether the subject looked to the target location again after checking back. 10 percent of the videos were coded for inter-observer reliability, and intraclass correlation was good (ICC = 0.85, F= 12.6, p<0.001).

### Statistical Analysis

The data were analysed with generalized linear mixed models (GLMMs) in RStudio (Version 1.4.1717) [67]. For every experiment, a model for each of the two conditions was created. The models were fitted with a binomial distribution, and individual identity of the observer was added as a random factor with session nested within. Head movements towards a target served as the response variable; species and trial types (luring stimulus present/not present), as well as their two-way interaction, were fixed factors. We reduced these full models, using the Akaike Information Criterion (AIC), to find those explaining the most variance. These final models were subjected to likelihood ratio tests to assess the effect of remaining factors (for values of final models, see Supplementary Material). If trial type with the lure stimulus present had no significant effect in the *no-demonstrator* condition, but a significant effect in *demonstrator* conditions, this was interpreted as gaze-following. Subsequently, we ran the same models for each experiment but used “checking-back” as the response variable. “Checking-back” behaviours were defined as first following the gaze and thereafter looking back to the demonstrator.

## Acknowledgements

This work was supported by Swedish Research Council grants 2019-03265 and 2021-02973. We thank Nina Thierij, Mark Kernkamp, Mathias Andersson, Morgan Luce, Simon Grendeus, Thomas Rejsenhus Jensen, Ola Oscarsson, and Ivo Jacobs for their practical help. We are grateful for Helena Osvath for the creation of figures. We thank Steve Brusatte, Pavel Nemec and Lawrence Witmer for helpful comments on the manuscript. We are also grateful to Ystad Zoo.

