## Supplementary Material for "Gaze following in Archosauria – Alligators and palaeognath birds suggest dinosaur origin of visual perspective taking"

#### 1. Ethical statement

All animals participated on voluntary basis and could leave the set-up, that was installed in a part of their home enclosure, whenever they chose to. No force of any kind was used. Sometimes animals were given food rewards. Both the observer and the demonstrator were familiar group members with no known antagonistic history. All animals were housed in either zoos or at private owners that met the legal requirements, as well as Lund Universities ethical standards. The research did not include so-called procedures under the EU-directive 2010/63/EU, and did not qualify for ethical approval, which is also true according to the stricter Swedish legislation (SJVFS 2019:9, chapter 2, § 22).

#### 2. Subjects & housing

Table 1: Test subjects

| Species | Subject | Sex | Age | Housing | Role |
|---|---|---|---|---|---|
| *A. mississippiensis* | Toke | female | subadult | group-housed | Demonstrator& Subject |
| *A. mississippiensis* | Ivar | male | subadult | group-housed | Demonstrator& Subject |
| *A. mississippiensis* | Bestla | male | subadult | group-housed | Subject |
| *A. mississippiensis* | Sigi | female | subadult | group-housed | Subject |
| *A. mississippiensis* | Kåra | female | subadult | group-housed | Subject |
| *A. mississippiensis* | Gudrun | female | subadult | group-housed | Subject |
| *D. novaehollandiae* | Snow | female | adult | pair-housed | Demonstrator& Subject |
| *D. novaehollandiae* | Tufty | male | adult | pair-housed | Demonstrator& Subject |
| *D. novaehollandiae* | Uncrowned | female | adult | pair-housed | Demonstrator& Subject |
| *D. novaehollandiae* | Crowned | male | adult | pair-housed | Demonstrator& Subject |
| *D. novaehollandiae* | Judy | female | adult | pair-housed | Demonstrator& Subject |
| *D. novaehollandiae* | Harry | male | adult | pair-housed | Demonstrator& Subject |
| *R. americana* | Nox | male | adult | group-housed | Demonstrator& Subject |

| | | | | | |
|---|---|---|---|---|---|
| *R. americana* | Hamilton | male | adult | group-housed | Demonstrator& Subject |
| *R. americana* | Yvette | female | adult | group-housed | Subject |
| *R. americana* | Salsa | female | adult | group-housed | Demonstrator& Subject |
| *R. americana* | Lucia | female | adult | group-housed | Demonstrator& Subject |
| *R. americana* | Arroz | female | adult | group-housed | Subject |
| *E. elegans* | Alicio | male | adult | group-housed | Demonstrator& Subject |
| *E. elegans* | Sleepy Genius | male | adult | group-housed | Demonstrator& Subject |
| *E. elegans* | Pretty Boy | male | adult | group-housed | Demonstrator& Subject |
| *E. elegans* | New Tinamou | male | adult | group-housed | Subject |
| *E. elegans* | Jon Snow | male | adult | group-housed | Subject |
| *E. elegans* | Sandy | female | adult | group-housed | Subject |
| *G. gallus* | Yellow | female | adult | group-housed | Demonstrator& Subject |
| *G. gallus* | Pink | female | adult | group-housed | Demonstrator& Subject |
| *G. gallus* | Red | female | adult | group-housed | Subject |
| *G. gallus* | White | female | adult | group-housed | Subject |
| *G. gallus* | Green | female | adult | group-housed | Subject |
| *G. gallus* | Rooster | male | adult | group-housed | Subject |

### 3. Housing and experimental procedures

#### *3.1. Alligators*

Subjects were six seven-year-old American alligators (*Alligator mississippiensis*; 2 males and 4 females) that were group-housed in an indoor facility consisting of a 42 $m^2$ pool area and a 26.5 $m^2$ land area. Subjects were tested on land by dividing the pool from the land area with opaque screens that didn't allow for visual contact with the rest of the group, but they could still hear each other.

The testing arena was in all experiments divided by a mesh barrier into two areas, one in which the demonstrator was placed (in the no-demonstrator condition, this compartment stayed empty) and one for the subject. Tests were conducted by three experimenters, one on the subject side, and two on the demonstrator side. The two experimenters on the demonstrator side were seated behind 60-centimeter-high wooden barriers on either side of the demonstrator from where they could present stimuli, position the animal, and throw food rewards (shrimps). The barrier provided security, but also prevented the animals from seeing arm movements of the experimenters that could have given away on which side the stimulus was presented. Both experimenters on the demonstrator side looked at the ground during trials, as not to give gaze cues themselves.

The general procedure of all experiments was as follows: The experimenters positioned both animals (or only the subject in the no-demonstrator condition) in front of the mesh divider. The experimenter on the subject side then stepped behind the animal and announced the beginning of the trial. Each trial began with a ten second baseline. Then, the test stimulus was presented, either until the demonstrator reacted by gazing in the indicated direction, or for 5 seconds in the no-demonstrator condition. After the demonstration, trials lasted for another 10 seconds.

*3.2. Small birds*

Small birds in this study were six adult elegant-crested tinamous (*Eudromia elegans*, 1 female and 5 males) and six adult red junglefowl (*Gallus gallus*, 1 male and 5 females). Elegant crested tinamous were group-housed in an outdoor aviary, while red junglefowl had access to an indoor and outdoor aviary. Elegant crested tinamous were tested in their outdoor aviary, while red junglefowl were tested indoors.

Small birds were tested by one experimenter. In the demonstrator condition, the experimenter lured both animals to a mesh divider by throwing small pieces of food (mealworms) until they were facing each other. In the no-demonstrator condition, only the subject was guided into position. Birds did not receive a baseline, as they would have moved away from the barrier during that time. Trials lasted for 10 seconds after demonstration.

*3.3. Large birds*

Large birds in this study were six adult emus (*Dromaius novaehollandiae*, 3 females and 3 males) and six adult greater rheas (*Rhea americana*, 2 males and 4 females). Emus were all pair-housed, while rheas lived in two mixed-sex groups. Both species were tested in their outdoor enclosures. Again, trials started without a baseline, and lasted for 10 seconds.

### 4. Statistical models

Table 2: Results of likelihood ratio test performed on the final Generalized Linear Mixed Models (lowest AIC). VCO = visual co-orientation; TA = turning around; CB = checking back.

| Model | Response variable | Distribution | Coefficient | Chisq | Df | P |
|---|---|---|---|---|---|---|
| Proportions gaze following into distance | VCO | Binomial | Experimental Condition | 9.71 | 1 | 0.0022** |
| | | | Species | 25.80 | 4 | <0.001*** |
| Experiment 1, birds, no-demonstrator condition | VCO | Binomial | Species | 3.19 | 3 | 0.36 |
| | | | Test Condition | 0.12 | 1 | 0.73 |
| Experiment 1, birds, demonstrator-condition | VCO | Binomial | Species | 4.47 | 3 | 0.21 |
| | | | Test Condition | 16.33 | 1 | <0.001*** |
| | | | Species*Test Condition | 4.40 | 3 | 0.22 |
| Experiment 1, alligators, no-demonstrator condition | TA | Binomial | Test Condition | 1.86 | 1 | 0.17 |
| Experiment 1, alligators, demonstrator condition | TA | Binomial | Test Condition | 5.77 | 1 | 0.01* |
| Experiment 2, birds, no- | VCO | Binomial | Test Condition | 3.61 | 1 | 0.05 |

| | | | | | | |
|---|---|---|---|---|---|---|
| demonstrator condition | | | | | | |
| Experiment 2, birds, demonstrator condition | VCO | Binomial | Species | 9.26 | 3 | 0.03* |
| | | | Test Condition | 14.73 | 1 | <0.001*** |
| | | | Species*Test Condition | 5.67 | 3 | 0.13 |
| Experiment 2, alligators, no-demonstrator condition | VCO | Binomial | Test Condition | 1.39 | 1 | 0.24 |
| Experiment 2, alligators, demonstrator condition | VCO | Binomial | Test Condition | 4.09 | 1 | 0.04* |
| Experiment 3, birds, no-demonstrator condition | VCO | Binomial | Species | 3.96 | 3 | 0.27 |
| | | | Test Condition | 3.44 | 1 | 0.06 |
| Experiment 3, birds, demonstrator condition | VCO | Binomial | Species | 4.88 | 3 | 0.18 |
| | | | Test Condition | 33.74 | 1 | <0.001*** |
| Checking back birds | CB | Binomial | Experimental Condition | 4.65 | 2 | 0.098 |
| | | | Species | 9.72 | 3 | 0.021* |
| | | | Experimental Condition*Species | 13.75 | 6 | 0.033* |
